## Supplemental Methods for "Functional characteristics and computational model of abundant hyperactive loci in the human genome"

### 1. Supplemental Methods

#### 1.1 Datasets

TF, histone modification ChIP-seq and ATAC-seq datasets for HepG2, K562, and H1 were batch-downloaded from the ENCODE Project (Davis *et al.*, 2018). For each ChIP-seq target of each cell line, if there were multiple datasets, the one with the latest date was selected (Table S1). The GRCh37/hg19 assembly was used as a reference genome throughout the study. In those cases when the ChIP-seq dataset was reported on GRCh38/hg38, the coordinates were converted to hg19 using liftOver. The phastCons (46 vertebrates), CpG islands, repeat elements, and GENCODE TSS annotations were obtained from the UCSC genome browser database (Kent *et al.*, 2002). Transcribed enhancer regions (eRNAs) were obtained from the FANTOM database (Lizio *et al.*, 2019). Super-enhancer regions were obtained from (Hnisz *et al.*, 2013). GC contents were calculated using the “nuc" functionality of the bedtools program (Quinlan and Hall, 2010). Gene expression data was obtained from the Roadmap Epigenomics project (https://egg2.wustl.edu/roadmap/data/byDataType/rna/expression/). Tissue-specificity metric *tau* scores for genes were downloaded from (Palmer *et al.*, 2021).

#### 1.2 Definitions

The loci were divided into bins according to a two-part scale. The first part is on a linear scale from 1 to 5, and the second part is on a logarithmic scale from 5 to the maximum number of DAPs bound to a single locus in that cell line.

| | **TFs** | **Bin edges (n=14)** | | | | | | | | | | | | | | |
|---|---|---|---|---|---|---|---|---|---|---|---|---|---|---|---|---|
| HepG2 | 545 | 1 | 2 | 3 | 4 | 5 | 7 | 12 | 19 | 31 | 48 | 77 | 122 | 192 | 304 | 480 |
| K562 | 411 | 1 | 2 | 3 | 4 | 5 | 7 | 11 | 16 | 24 | 37 | 55 | 82 | 123 | 184 | 275 |
| H1 | 47 | 1 | 2 | 3 | 4 | 5 | 6 | 7 | 8 | 10 | 12 | 15 | 18 | 22 | 26 | 32 |
| | | **linear growth (n=4)** | | | | | **logarithmic growth (n=10)** | | | | | | | | | |

These nominal numbers are used in cases when the distributions are displayed for individual cell lines (eg. Fig1A). When the figures display the distributions for two cell lines in a joint manner (eg Fig3A), the edges are converted to the average percentages of the overall scale lengths for each cell line. HOT regions are defined as loci with the number of DAPs corresponding to the last four bins.

*Regular enhancers* were defined as the central 400bp regions of DHS which overlap H3K27ac histone modification regions with promoter and exons removed from them.
*Promoters* were defined as 1.5kbs upstream and 500 downstream regions of the canonical and alternative TSS coordinates, extracted from the *knownGenes.txt* table obtained from UCSC Genome Browser database. All the genomic arithmetic operations were done using the *bedtools* program. Figures were generated using Matplotlib (Hunter, 2007) and Seaborn (Waskom, 2021) packages. Statistical and numerical analyses were done using the *pandas*, *NumPy*, *SciPy,* and *sklearn* packages (Virtanen *et al.*, 2020). Genomic repeat regions were

obtained from http://www.repeatmasker.org/. CpG islands were extracted from *cpgIslandExt* table obtained from the UCSC Genome Browser.

#### 1.3 Joint DAPs and Hi-C 3D chromatin analysis.

To jointly analyze the conditional distributions of ChIP-seq signal levels in presence/absence of individua DAPs, we extracted a square matrix of size n=545 using the DAPs present in HepG2. For each analyzed 400bp locus, we extracted bound DAPs together with the ChIP-seq signal values. Then, each binary combination of the DAPs bound in that locus is added to their respective cells on the square matrix. Afterward, the matrix is normalized along the x-axis with the maximum value of each row. In other words, each row represents the normalized ChIP-seq signal strength in the presence of DAP indicated on the x-axis. We treated the empty values as 0 and removed the main diagonal values, leading to the matrix size of 544x544. Hierarchical clustering was done using the UPGMA algorithm.

Hi-C data analysis was carried out on HepG2. The datasets were obtained from ENCODE Project: ENCFF050EKS (chromatin loops), ENCFF018XKF (TADs), ENCFF548XLR (hic file). The coordinates in all of the datasets were converted from hg38 to hg19 using LiftOver.

The significant long-range contacts with 5 kbs resolution were extracted using the FitHiChIP program (Bhattacharyya *et al.*, 2019) with a threshold q-value<0.0001 and ICE bias correction, using *all-against-all* option.

The loci with >50% overlap were considered for the analysis of loops, TADs, and long-range chromatin contacts. Using the long-range chromatin contacts, we constructed a graph such that each node is the analyzed 400 bp locus and the edge is a long-range chromatin contact if the two connected nodes are located on different legs of the chromatin contacts. Based on this graph, we calculated the total number of contacts between the loci located in different bins, leading to a 14x14 matrix. We then normalized the values in each cell of the matrix with the maximum number of contacts in all cells.

FIRE loci were extracted from the *.hic* file using FIRECaller R package (Schmitt *et al.*, 2016).

For enrichment analyses of all the mentioned Hi-C-related regions, the ATAC-seq regions were used as background.

#### 1.4 PPI enrichment analysis.

To test the significance of the PPI networks described above, we ran 100 trials for each cluster by randomly selecting an equal number of DAPs reported in PPI networks and calculated the significance of the PPI enrichment p-values. All of the reported PPI enrichment p-values were significantly higher than the randomized trials (p-value < 0.01, one-sample t-test).

PPI networks and PPI enrichment p-values were extracted using the STRING Database's API (https://string-db.org/cgi/help.pl?subpage=api). For each cluster of DAPs analyzed, we submitted the list of DAPs as identifiers and retrieved the p-values using the *ppi_enrichment* interface. For each cluster, we extracted 100 PPI enrichment p-values each time randomly selecting DAPs in equal numbers to the size of the analyzed cluster. We then used the set of 100 p-values as a background distribution and conducted a one-sample t-test, where by the null hypothesis the p-value of the cluster is the mean of 100 p-values and computed the p-values of significance of the reported PPI network. The results of this analysis are in Table S2.

#### 1.5 Statistical analyses

All the statistical significance analyses were done using the *SciPy* package. The p-values too small to be represented by the command line output were represented as <1E-100. Correlation values with the number of bound DAPs were calculated using the average of the value for the bins, and the midpoint numbers of the edges of each bin.

#### 1.6 Classification analyses

The aim of this section is to determine whether the HOT loci can be accurately predicted based on their DNA sequences alone, and sequence features, including GC, CpG, GpC contents, and CpG island coverage. For sequence-based classification, we trained a Convolutional Neural Network (CNN) model using one-hot encoded sequences and an SVM classifier trained on gapped k-mers (gkmSVM) (Supplemental Methods 1.6.2, Figure S12A). For feature-based classification, we trained logistic regression (LogReg) classifiers and SVM classifiers with linear, polynomial, radial basis functions and sigmoid kernels. We carried out the classification experiments using the following control sets: a) randomly selected loci from merged DNaseI Hypersensitivity Sites (DHS) of cell lines in the Roadmap Epigenomics Project, b) promoter regions, and c) regular enhancers.

Using the sequence features, we trained separate models using each of the features in addition to one with all of the features combined. We observed that, when averaged across all the methods, GC content value possesses the highest amount of discrimination power (auROC: 0.73), followed by the combination of all features (auROC: 0.70) (Figure S13A,B). When compared across the classification methods, LogReg and SVM with linear kernel outperformed the other non-linear kernels by 20%, suggesting that the features possess linearly combined or largely overlapping effects in encoding the information in HOT loci (Figure S13A).

When classified using the sequences directly, CNN yielded the highest performance with auROC of 0.91, while for the gkmSVM it was 0.86 (both averaged over cell lines and control sets), suggesting that CNNs capture the motif grammar of the HOT loci better than gapped k-mers (Figure S13C). When the two classification schemes (sequence- and feature-based) are compared, CNNs outperformed the LogReg and linear SVMs by a factor of 1.3x (or 17%), suggesting that there is additional information that is highly relevant to the DNA-DAP interaction density encoded in the DNA sequences, in addition to the GC, CpG, GpC (Fig 5C).

##### 1.6.1 Datasets

For classification experiments of different categories of regions, the loci from HepG2 and K562 were used. For the classification of HOT loci, three different setups were constructed using the control (negative) sets:

- Randomly selected from the merged DHS regions obtained from the Roadmap Epigenomics Project to be 10x the size of the positive set (HOT loci)
- Regular enhancers (see 1.2), with the HOT loci subtracted
- Regular promoters (see 1.2)

The regions from chromosomes 6,7 were used as validation sets, the chromosomes 8, and 9 were used as test sets, and the rest of the autosomal chromosomes were used as training sets.

The total number of regions in classification setups and their train/validation/test sets splits is as follows:

| Controls: DHS | | | | | |
|---|---|---|---|---|---|
| Cell Line | HOTs | Controls | Train | Validation | Test |
| HepG2 | 25,928 | 249,499 | 210,520 | 33,231 | 31,676 |
| K562 | 15,231 | 146,585 | 123,041 | 20,310 | 18,465 |
| Controls: regular enhancers | | | | | |
| Cell Line | HOTs | Controls | Train | Validation | Test |
| HepG2 | 25,928 | 249,499 | 210,520 | 33,231 | 31,676 |
| K562 | 15,231 | 146,585 | 123,041 | 20,310 | 18,465 |
| Controls: regular promoters | | | | | |
| Cell Line | HOTs | Controls | Train | Validation | Test |
| HepG2 | 25,928 | 28,621 | 34,970 | 5,479 | 3,403 |
| K562 | 15,231 | 25,810 | 41,979 | 5,800 | 3,959 |

#### 1.6.2 Sequence-based classification

##### 1.6.2.1 Convolutional Neural Networks (CNNs)

For training CNNs, the sequences of the loci were converted to one-hot encoding, with the lengths options of 400bps and extended to 1000bps.
The model consists of the layers as follows.

| Layer | Params | Activation |
|---|---|---|
| 1.Convolutional | filters=480, kernel_size=9, stride=1 | ReLu |
| 2.Max pool | Pool_size=9, stride=3 | |
| 3.Droupout | P=0.2 | |
| 4.Convolutional | filters=480, kernel_size=4, stride=1 | ReLu |
| 5.Max pool | Pool_size=4, stride=2 | |
| 6.Droupout | P=0.2 | |
| 7.Convolutional | filters=240, kernel_size=4, stride=1 | ReLu |
| 8.Max pool | Pool_size=4, stride=2 | |
| 9.Droupout | P=0.2 | |
| 10.Convolutional | filters=320, kernel_size=4, stride=1 | ReLu |
| 11.Max pool | Pool_size=4, stride=2 | |
| 12.Fully connected | units=180 | ReLu |
| 13. Fully connected | units=15 | Sigmoid |

Total number of trainable parameters is 2,342,723. The kernels were subjected to constraints of max_norm=0.9, l1=5*10E-7, l2=1E-8. Each instance of the model was trained using the input lengths of 400 and 1000. The training process was optimized using the Adadelta optimizer (“[1212.5701] ADADELTA: An Adaptive Learning Rate Method,” n.d.) and ran for a maximum of 200 epochs with a patience period of 15. The models were built using *tensorflow v2.3.1* and trained on NVIDIA k80 GPUs.

##### 1.6.2.2 SVM classification.

For SVM classification, gapped k-mer SVM program was used downloaded from https://github.com/Dongwon-Lee/lsgkm. For each category of the regions, instances of SVM models were trained, using 400bp and 1000bp regions, with the following kernel options:

*0 -- gapped-kmer*
*1 -- estimated l-mer with full filter*
*2 -- estimated l-mer with truncated filter (gkm)*
*3 -- gkm + RBF (gkmrbf)*
*4 -- gkm + center weighted (wgkm)*
*5 -- gkm + center weighted + RBF (wgkmrbf)*

Of the tried kernels, gapped-kmer kernel outperformed the others, so the gapped-kmer option was used for further comparison with other methods.

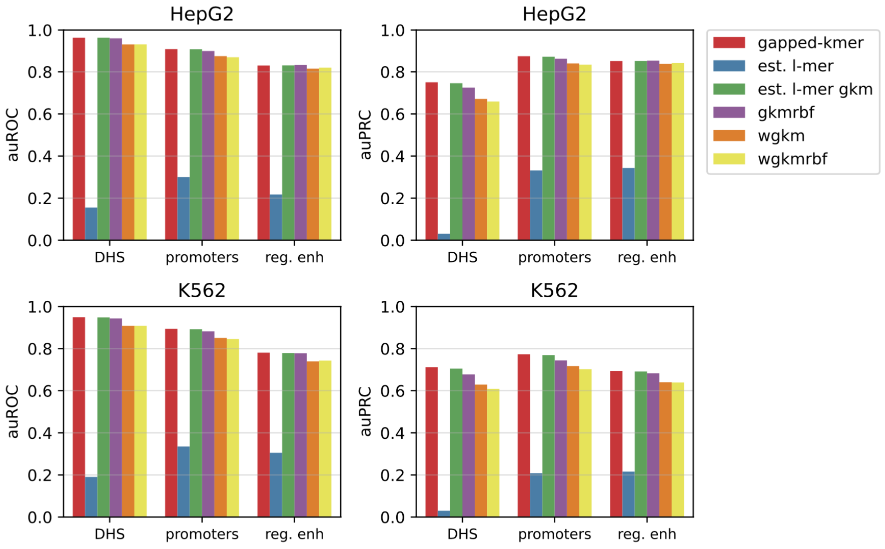

SVM performed worse with 1000kbs input length compared to 400 bp. So, 400bp results were used for further comparisons with other methods.

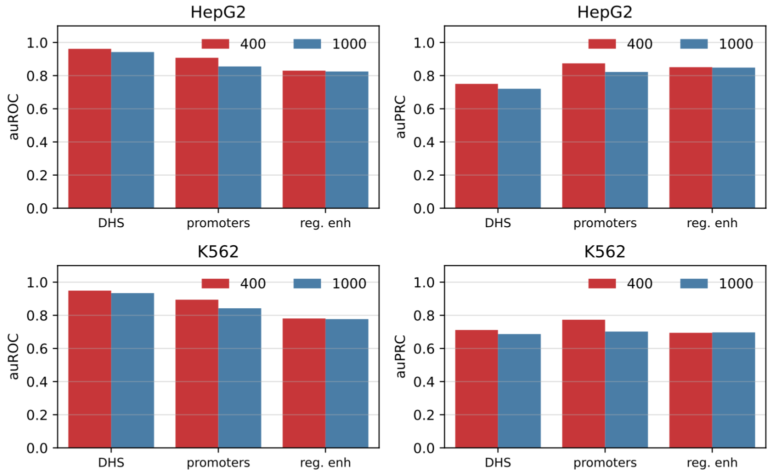

#### 1.6.3 Feature-based classification

The features used for classification were:
- GC content
- CpG content: counted the occurrences of “CG“ as density over the sequence length.
- GpC content: counted the occurrences of “GC“ as density over the sequence length.

- CpG island coverage: fraction of the overlaps with the CpG island obtained from UCSC Genome Browser database.

Each classification model was trained using all of the features at once (n=4), and using each of the features separately.

Logistic regression:
*sklearn.linear_model.LogisticRegression* method was used from scikit-learn library.

SVM:
sklearn.svm.SVM method was used from scikit-learn library

Kernels used with SVM classification are: *linear*, *polynomial*, *radial basis function(rbf)*, and *sigmoid*.

#### 1.7 Variant analysis

Common SNPs and INDELs were extracted from the *gnomAD r2.1.1* dataset (Karczewski et al. 2020). Variants with PASS filter value and MAF>5% were selected using the "*view -f PASS -i 'MAF[0]>0.05'*" options of *bcftools* program (Li, 2011). raQTLs were downloaded from https://sure.nki.nl (van Arensbergen et al., 2019). Liver and blood eQTLs were extracted from the GTEx v8 dataset (https://www.gtexportal.org/home/datasets). Liver caQTLs were obtained from the supplementary material of (Currin et al., 2021). NHGRI-EBI GWAS database variants were grouped according to their traits (dataset e0_r2022-11-29). For each GWAS SNP, LD SNPs with r2>0.8 were added using the *plink v1.9* (Chang et al., 2015) program using the parameters "*--ld-window-r2 0.8 --ld-window-kb 100 --ld-window 1000000*". Enrichments of GWAS-trait SNPs were calculated as the ratios of densities of SNPs in each class of regions (eg. HOT enhancers, HOT promoters) to either that of the regular enhancers or the DHS regions. The statistical significance of enrichment was calculated using the binomial test. FDR values were calculated using the Bonferroni correction.

### 2. Supplemental Figures

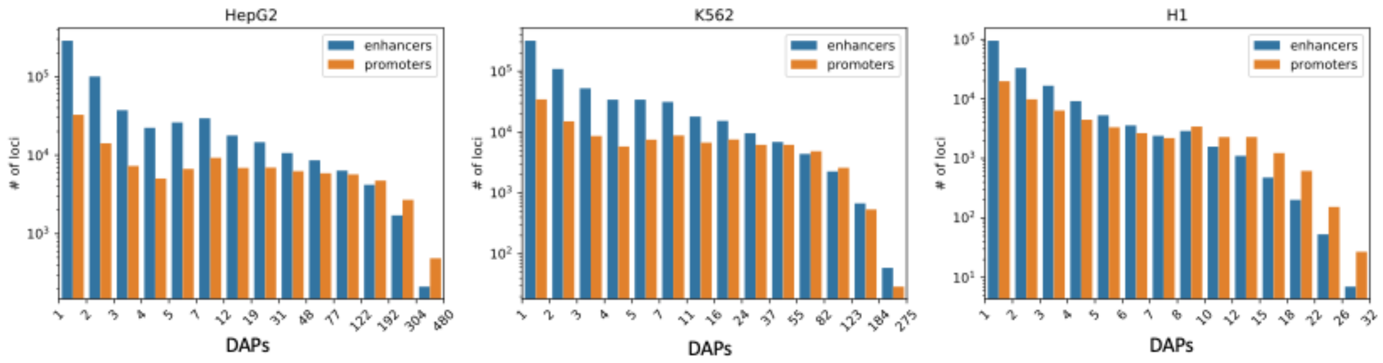

**Fig S1**. Distribution of the number of loci by the number of overlapping peaks 400bp loci in HepG2, K562, and H1. Loci are binned on a logarithmic scale (Table 1. see Methods). Shaded region represents the HOT loci.

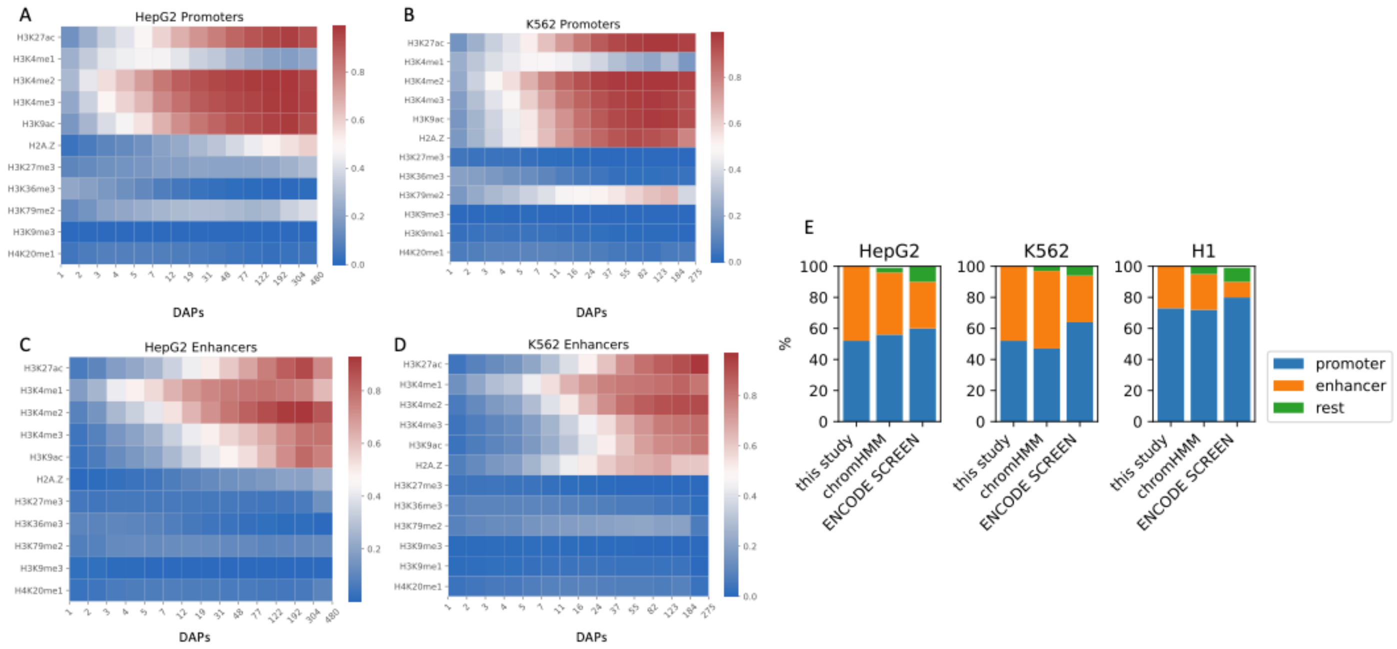

**Fig S2.** A, B) Percentages of overlapping promoter loci binned by bound DAPs with histone modification regions. C, D) Percentages of overlapping enhancer loci (non-promoter) binned by bound DAPs with histone modification regions. E) Composition of the HOT loci to promoter and enhancer regions based on the definitions used in this study, chromHMM states and ENCODE SCREEN annotations.

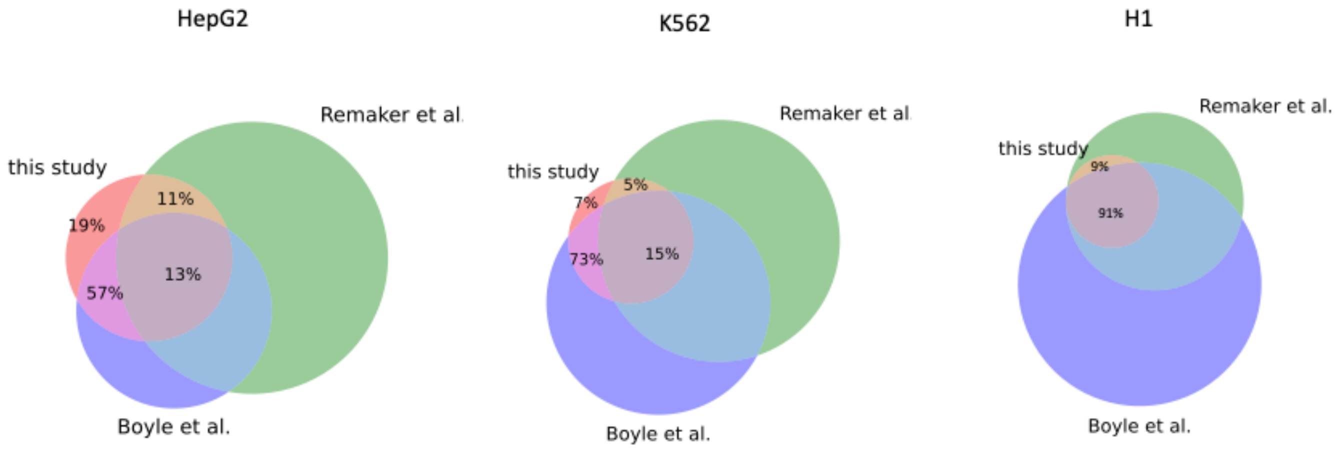

**Fig S3.** Overlaps between the HOT loci as reported in this study, Remaker et al. and Boyle et al. Overlaps are calculated in terms of fractions of overlapping bps.

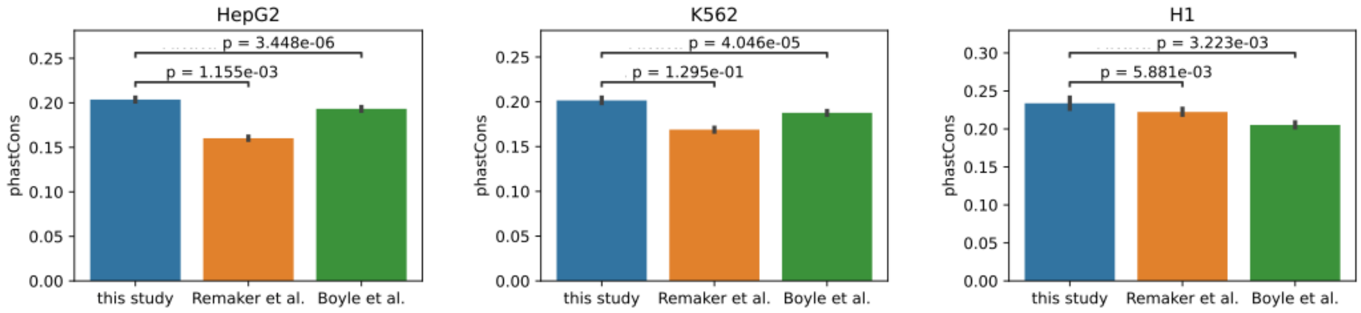

**Fig S4**. phastCons conservation scores of HOT loci defined by this study, Remaker et al., and Boyle et al. Bar plots depict median values, error bars are 95% confidence intervals. P-values are Mann-Whitney U test results.

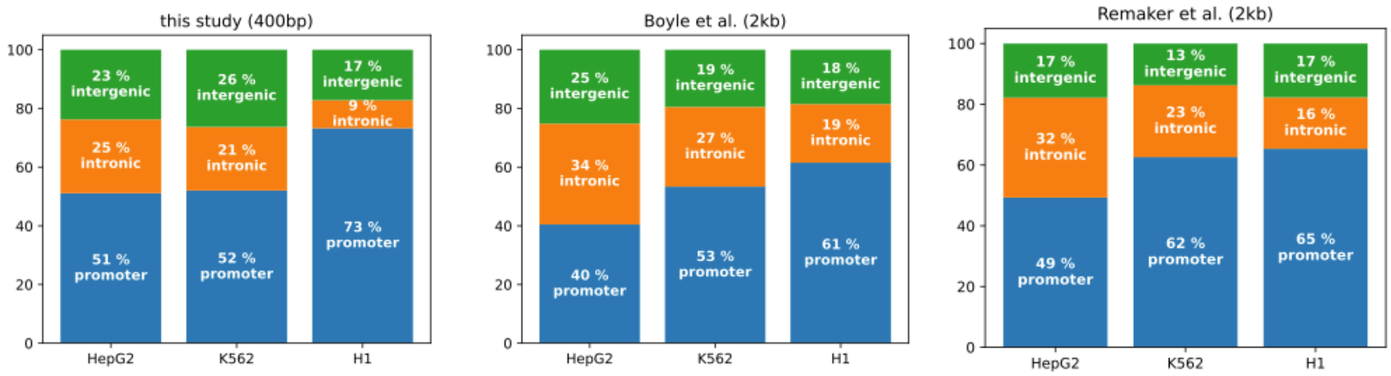

**Fig S5**. Compositions of HOT loci as reported in this study, Remaker et al. and Boyle et al. in terms of promoter, intronic and intergenic regions.

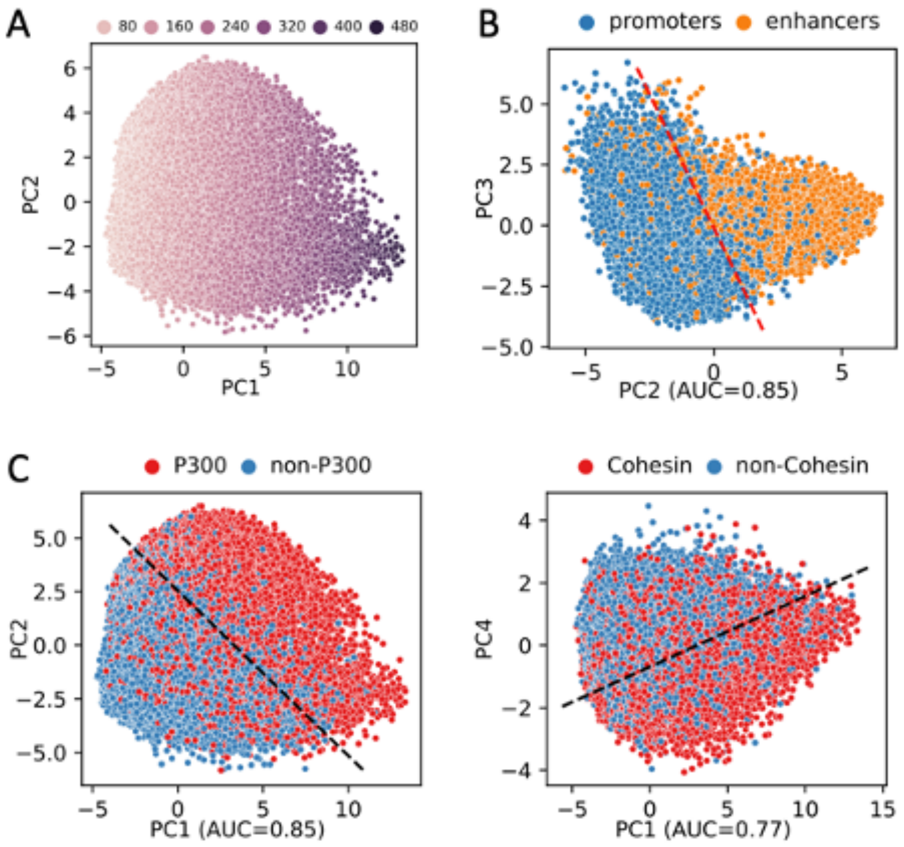

**Fig S6** PCA plots of HOT loci in HepG2 based on the DAP presence vectors. Each dot represents a HOT locus: **A**) PC1 and PC2 correlated with the number of overlapping DAPs. **B**) PC2 and PC3, with promoter and enhancer marked. **C**) PC1 and PC2, marked p300 bound HOT loci. **D**) PC1 and PC4, marked Cohesin bound HOT loci.

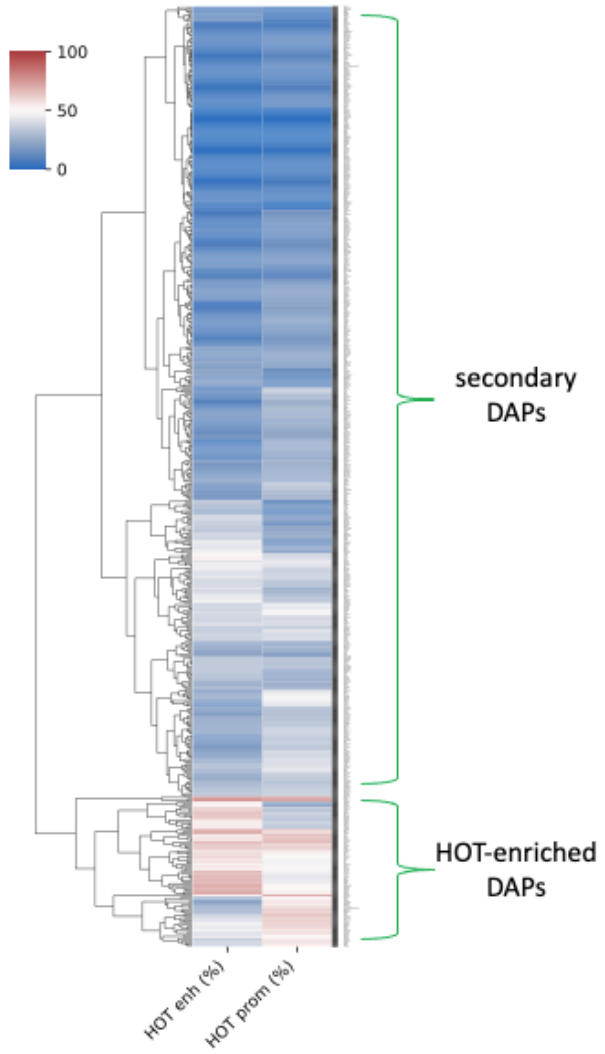

**Fig S7**. DAPs clustered by percentage of HOT promoters and HOT enhancers that the ChIP-seq peaks overlap. The top cluster comprises the DAPs which on average overlap 13% of HOT loci. The DAPs which form the bottom cluster are present in 53% of HOT loci.

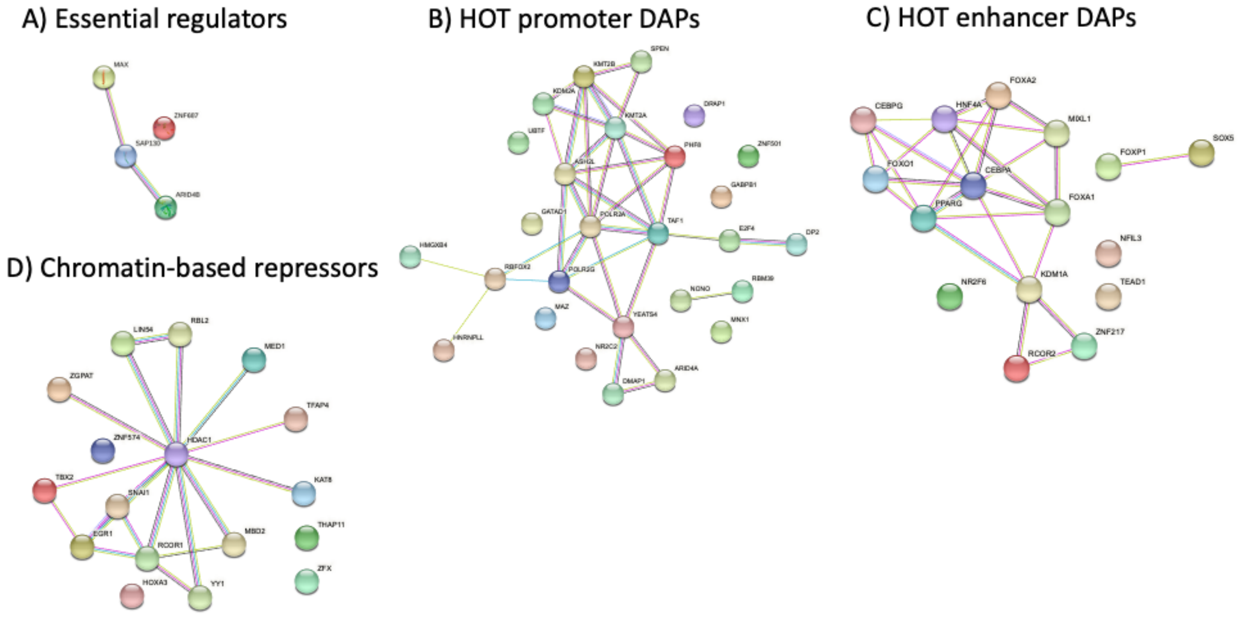

**Fig S8**. PPI networks of 4 clusters. Names of the clusters are indicated as titles. Refer to the text for interpretations.

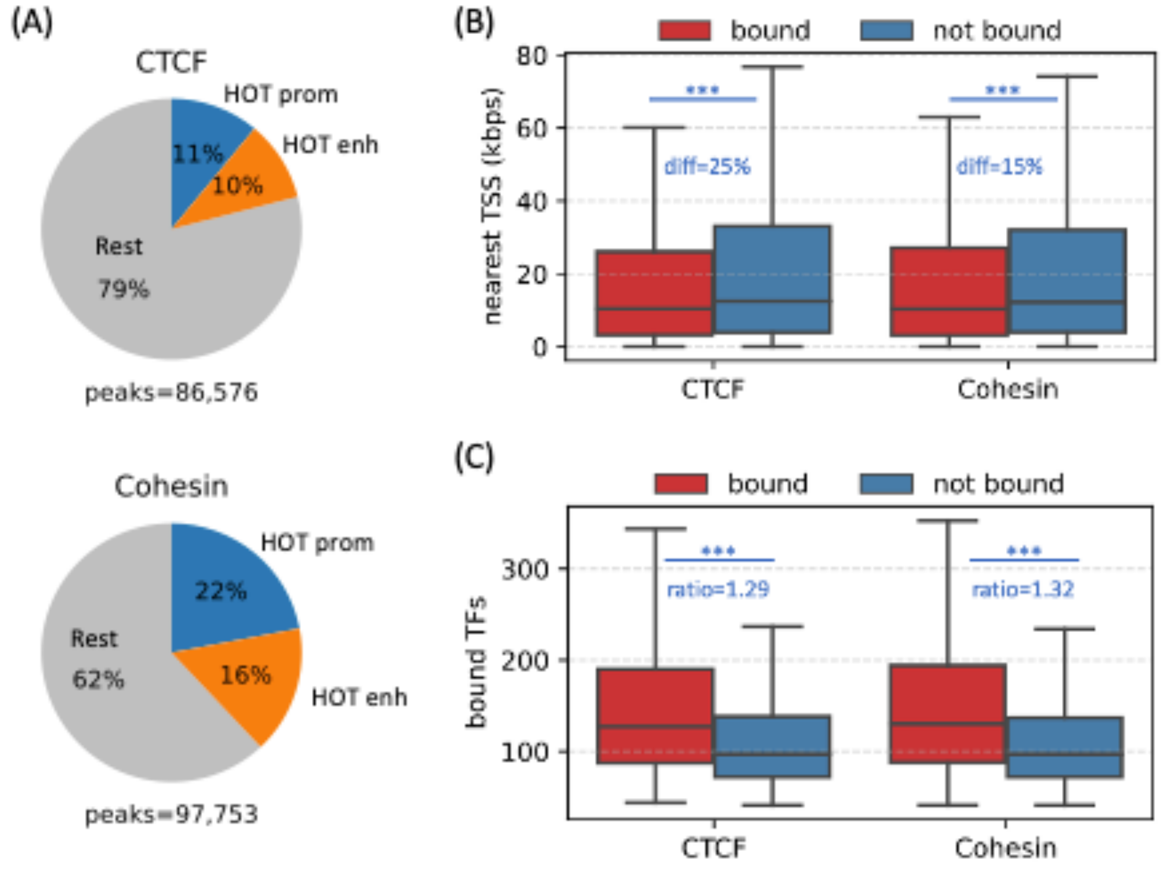

**Fig S9** CTCF and Cohesin in HOT loci. **A**) Fractions of CTCF and Cohesin ChIP-seq peaks in HOT promoters and enhancers. **B**) Distances to the nearest TSSs in HOT loci bound by CTCF and Cohesin. **C**) Numbers of total DAPs in HOT loci bound by CTCF and Cohesin.

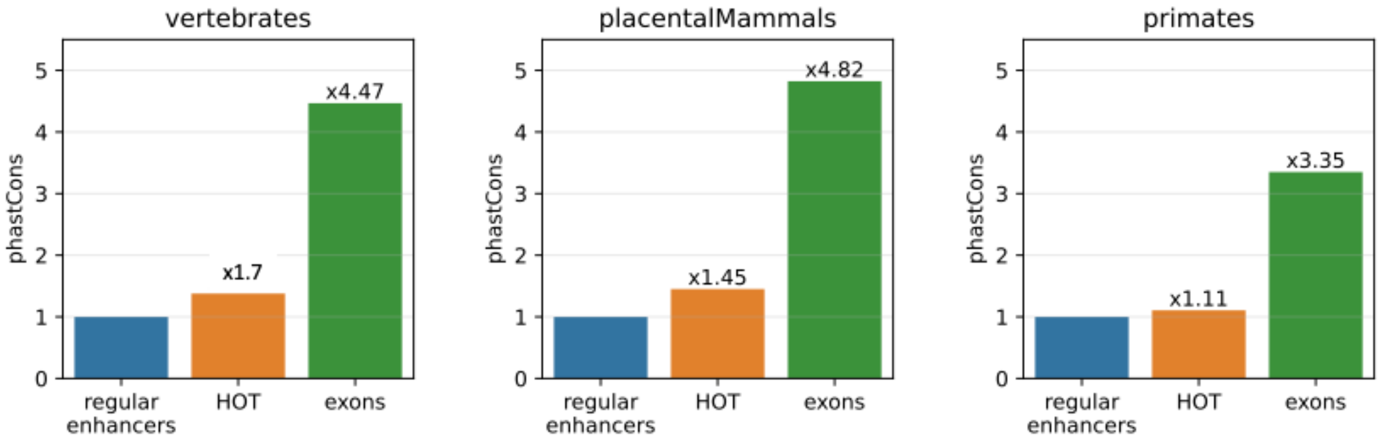

**Figure S10.** Comparison of phastCons conservation scores of regular enhancers, HOT loci and exons using the score extracted from vertebrates, placental mammals and primates.

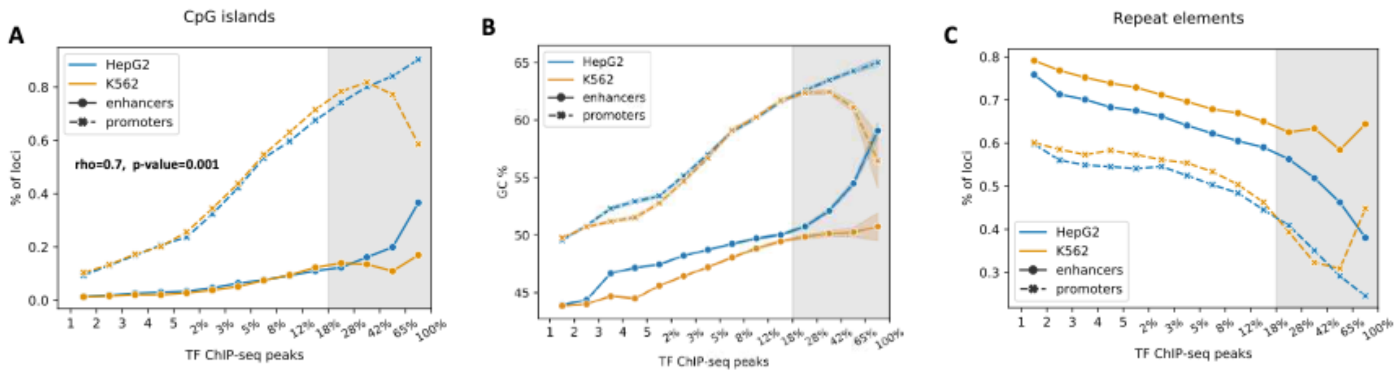

**Figure S11.** Sequence features of HOT loci. **A**) fractions of DAP-bound loci overlapping CpG islands. X-axis is bins of number of bound DAPs. The logarithmic bins are represented in terms of percent of total number of DAPs in given cell line. **B**) GC contents of DAP-bound loci. X-axis is the same as in **A**. **C**) Fractions of loci DAP-bound overlapping repeat elements. X-axis is the same as in **A**.

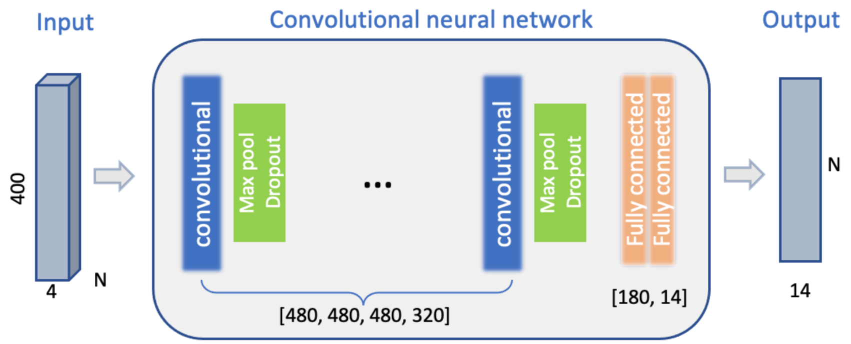

**Figure S12.** Schematic representation of trained CNN model.

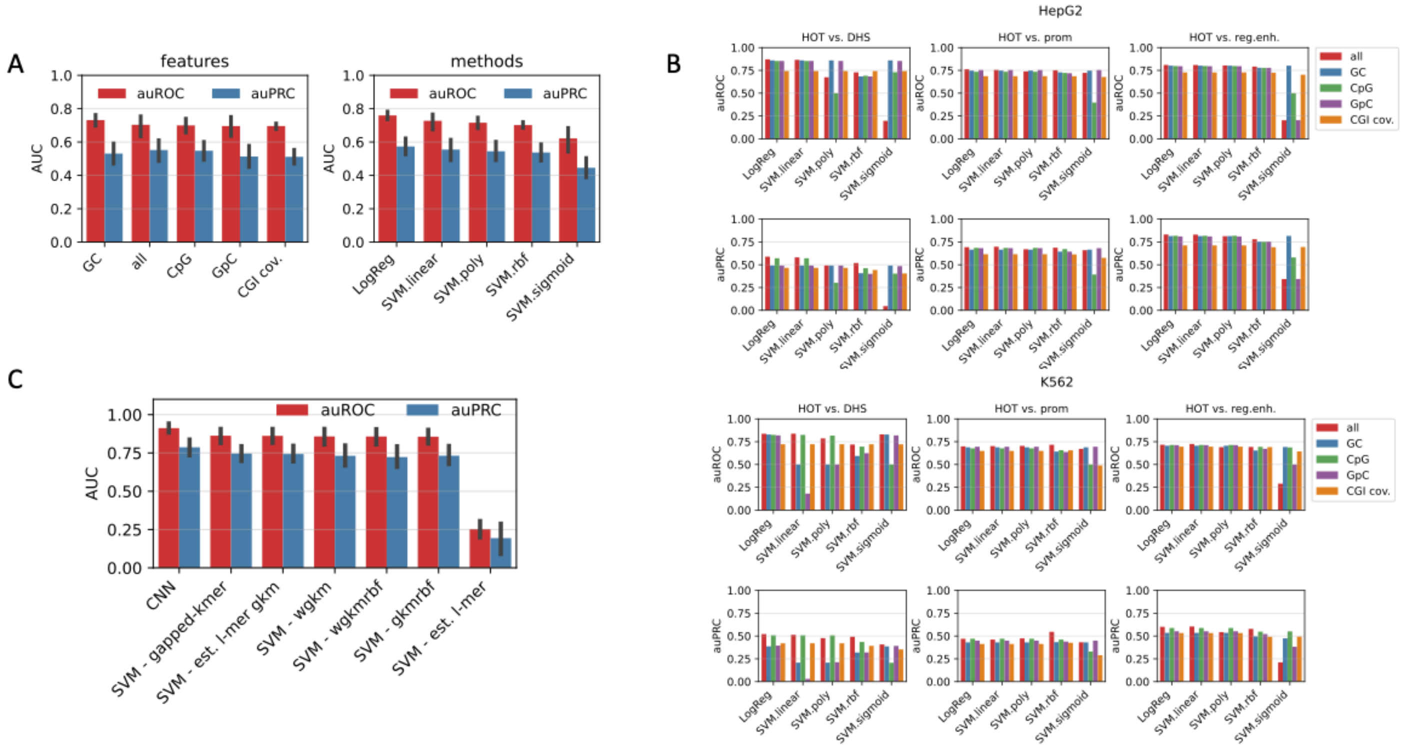

**Figure S13.** A) (*Left*) Average classification auROC/auPRC values of used sequence features. *(Right)* Average classification values of methods run on sequence features. B) Breakdown of auROC/auPRC values for each feature/method/background. C) Average classification metric values of sequence-based methods: CNNs and SVM with 6 different kernels.

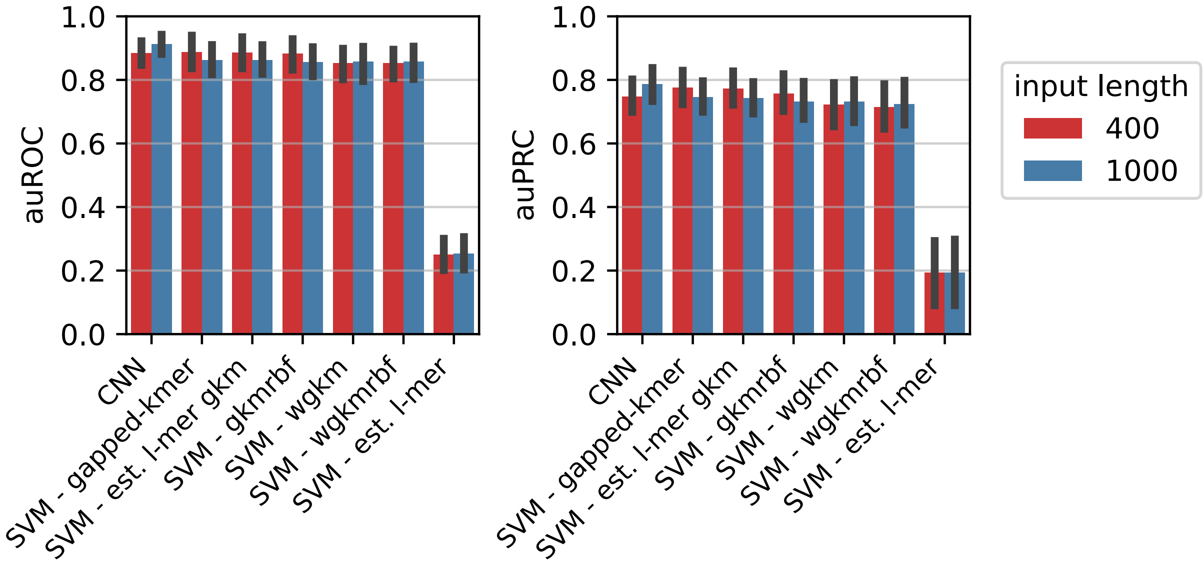

**Figure S14.** Comparison of classification performances for sequences in different lengths.

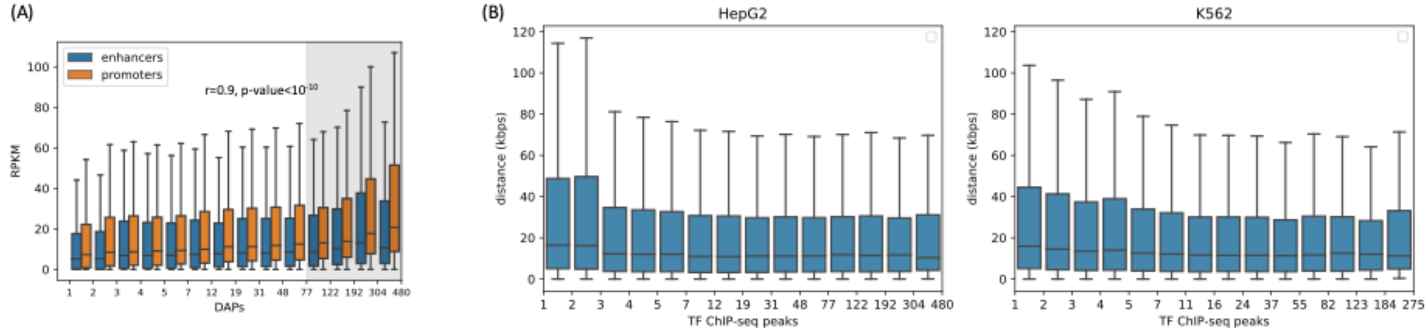

**Figure S15.** A) Expression levels of target genes of DAP-bound loci in HepG2. B) Distance to the nearest TSS from the DAP-bound non-promoter loci in HepG2 and K562.

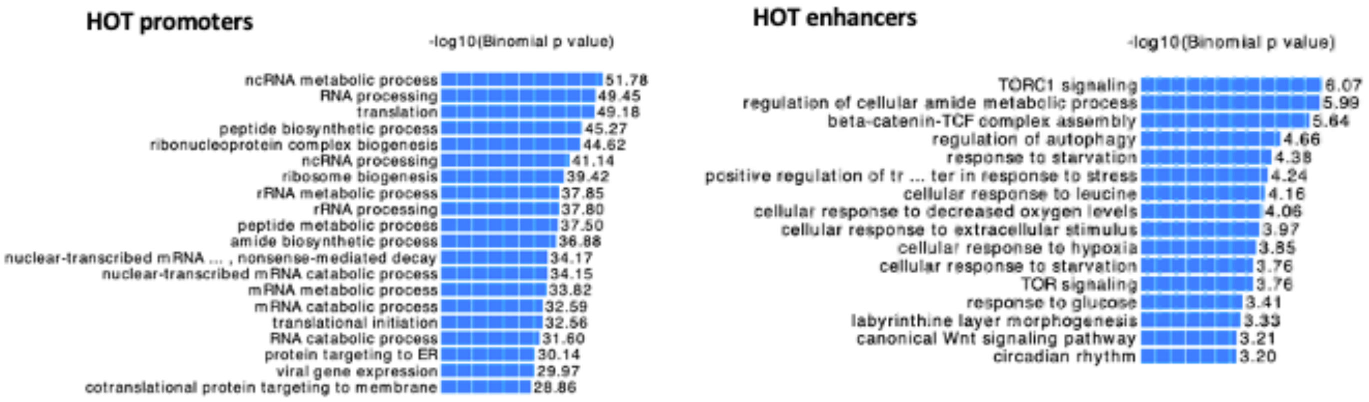

**Figure S16.** GO terms associated with the HOT enhancers and promoters in H1-hESC.

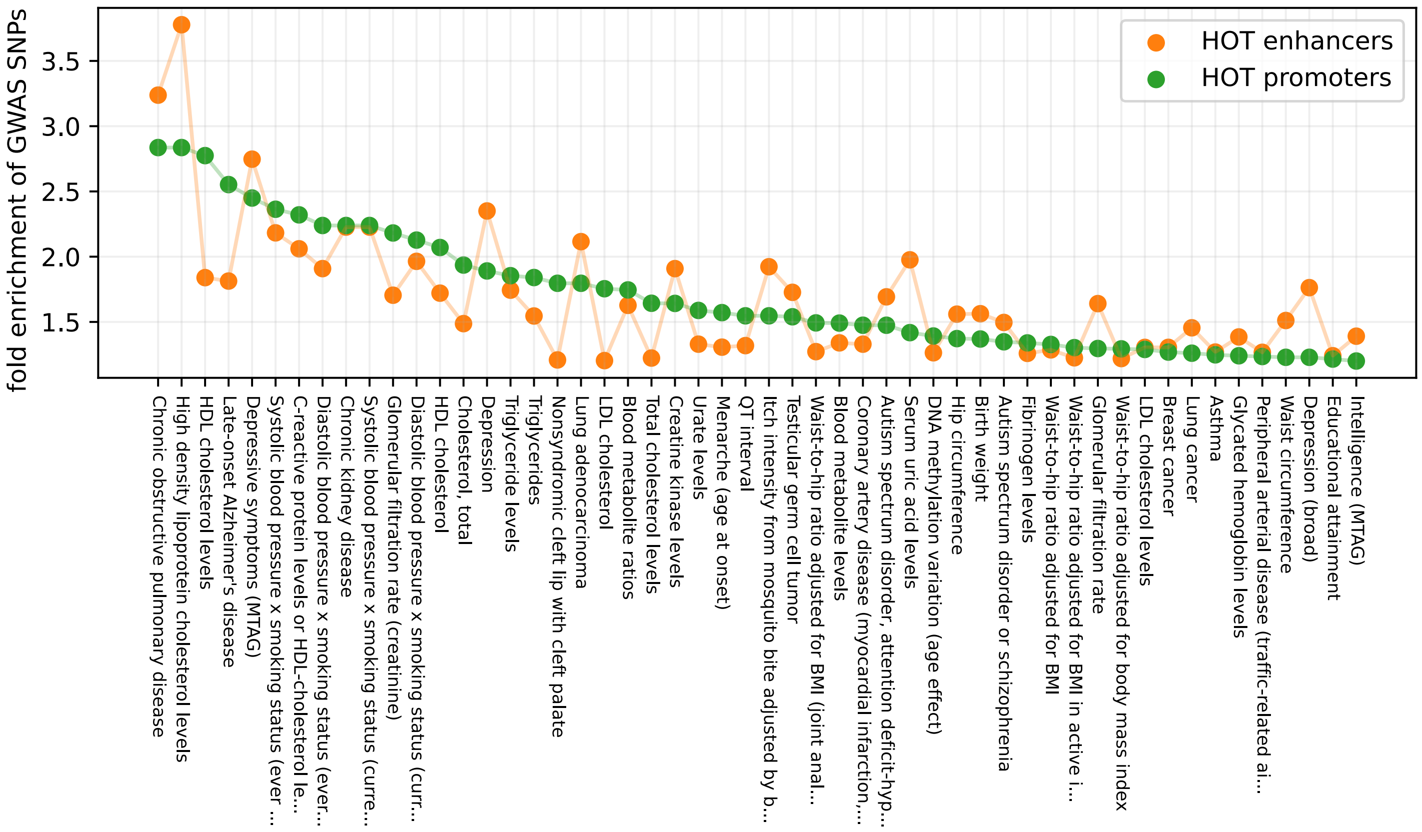

**Figure S17.** GWAS traits enrichment analysis filtered by unadjusted p-values (p-value<0.001).

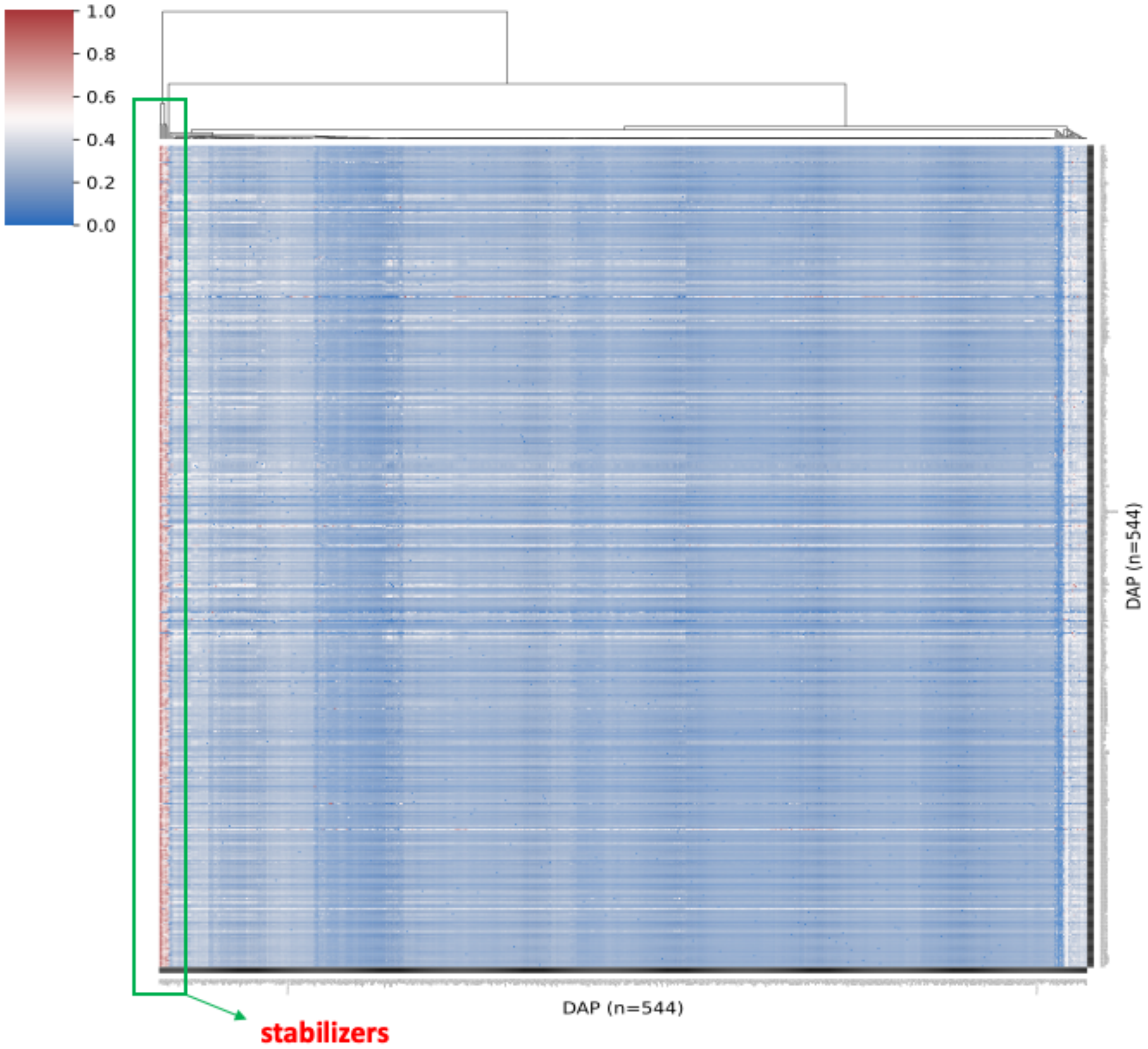

**Figure S18.** Normalized ChIP-seq signal values of DAPs in HOT loci (rows) in the presence of other DAPs (columns). The hierarchical clustering is done using the columns. That is, the leftmost outer group (in the green box) contains the DAPs in the presence of which most of the other DAPs yield highest ChIP-seq signal values.

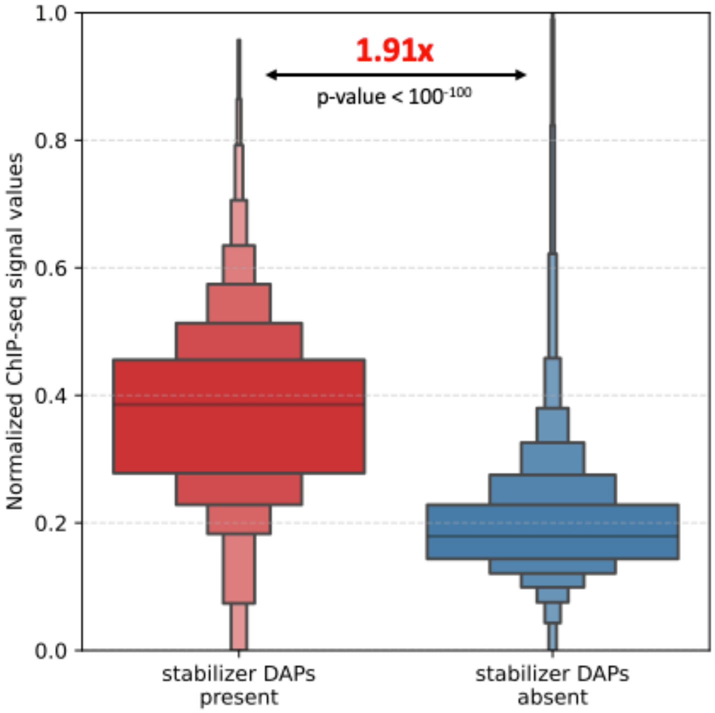

**Figure S19.** Distribution of the ChIP-seq signal values of DAPs when the stabilizing DAPs (Figure S18) are present vs. absent.

Palmer D, Fabris F, Doherty A, Freitas AA, de Magalhães JP. Ageing transcriptome meta-analysis reveals similarities and differences between key mammalian tissues. Aging (Albany

NY) 2021; 13: 3313–41.

Quinlan AR, Hall IM. BEDTools: a flexible suite of utilities for comparing genomic features. Bioinformatics 2010; 26: 841–2.

Schmitt AD, Hu M, Jung I, Xu Z, Qiu Y, Tan CL, et al. A compendium of chromatin contact maps reveals spatially active regions in the human genome. Cell Rep 2016; 17: 2042–59.

Virtanen P, Gommers R, Oliphant TE, Haberland M, Reddy T, Cournapeau D, et al. SciPy 1.0: fundamental algorithms for scientific computing in Python. Nat Methods 2020; 17: 261–72.

Waskom M. seaborn: statistical data visualization. JOSS 2021; 6: 3021.

[1212.5701] ADADELTA: An Adaptive Learning Rate Method [Internet]. [cited 2020 Oct 28] Available from: https://arxiv.org/abs/1212.5701
